# Computational evaluation of light propagation in cylindrical bioreactors for optogenetic mammalian cell cultures

**DOI:** 10.1101/2023.02.01.526707

**Authors:** Shiaki A. Minami, Priya S. Shah

## Abstract

Optogenetic control of cellular pathways and gene circuits in mammalian cells is a new frontier in mammalian genetic engineering. As a low-cost, tunable, and reversible input, light is highly adept at spatiotemporal, orthogonal regulation of cellular behavior. However, light is absorbed and scattered as it travels through media and cells, and the applicability of optogenetics in larger mammalian bioreactors has not been determined. In this work, we computationally explore the size limit to which optogenetics can be applied in cylindrical bioreactors at relevant height-to-diameter ratios for mammalian cell culture. We model the propagation of light using the radiative transfer equation and consider changes in reactor volume, absorption coefficient, scattering coefficient, and scattering anisotropy. We observed sufficient light penetration for activation for bioreactor sizes of up to 80,000 L with maximal cell densities, with decreasing efficiency for larger bioreactors. For a 100,000 L bioreactor, we determined that lower cell densities of up to 1.5·10^7^ cells/mL can be supported. We conclude that optogenetics can be applied to bioreactors at an industrial scale and may be a valuable tool for specific biomanufacturing applications.

## Introduction

Electromagnetic radiation in the visible range regulates cellular behavior in organisms that utilize light, such as plants, algae, and bacteria, as well as eyes in animals. The responsiveness to light is caused by opsins, which absorb photon energy and undergo conformational changes^[1]^. In optogenetics and other light-inducible synthetic biology systems, opsins that homo/hetero-dimerize, oligomerize, or undergo conformational changes upon light activation are used to control gene expression or protein activity. Optogenetics has enabled advancements in a variety of fields^[2,3]^, such as regulating and understanding cell motility in cell biology^[4]^, regulating specific pathways such as autophagy in cellular engineering^[5]^, or restoring vision in a blind patient in a biomedical application^[6]^. The spatiotemporal control and ease of use that optogenetics offers are clear advantages that have been demonstrated at a small scale in mammalian cell culture.

While the applications of optogenetics continue to expand, industrial applications will rely on bioreactors capable of transmitting light, or photobioreactors. Yet, as light travels through media and cells, its intensity decreases through absorption and scattering. Thus, understanding the design constraints of such photobioreactors is critical to establishing the feasibility of optogenetic approaches in industrial applications. In other bioproduction systems such as algae and cyanobacteria, cellular growth in photobioreactors has been studied, taking into consideration the attenuation of light in the bioreactor through absorption and scattering^[7–10]^. On small scales, optogenetics in model organisms such as *Saccharomyces cerevisiae* and *Escherichia coli* have also been performed on milliliter and liter scales to increase production^[11–13]^. However, propagation of light in mammalian bioreactors has not yet been evaluated. Given that reactor geometry heavily differs between microbial and mammalian bioreactors, it is critical to explore light transmission specific to typical mammalian bioreactor designs. Microbial photobioreactors often employ flat panel or tubular geometries to overcome limitations in light propagation, but mammalian cells often require cylindrical bioreactors with a limited range of tolerated aspect ratios for optimal cell growth or bioproduction^[14–16]^. If optogenetics is compatible with such reactor design, this technology may be useful for new biomanufacturing areas in which reactor design is still ongoing, including cultivated meat. This field is also constrained by considerably lower cost targets. Light-inducible systems could be far more economical than chemically-induced systems, which have added costs of the chemicals and potentially higher media costs if media exchange is required to remove the chemicals ^[15,17,18]^.

Here, we use computer simulations to explore light propagation in bioreactors with parameters tuned to mammalian cultures and bioreactor geometry. We find numerical solutions to the radiative transfer equation (RTE) and assess relevant ranges of absorption coefficients, scattering coefficients, scattering anisotropy, and reactor height-to-diameter ratios for mammalian cells. We determine that optogenetic systems are applicable to industrial scale bioreactors with high efficiencies up to 80,000 L.

## Materials and Methods

### Simulating radiation transfer in absorbing and scattering media

Computations were performed using COMSOL Multiphysics 5.5. Light propagation was modeled using Radiation in Absorbing Scattering Medium (RASM) physics in 3D. To model the propagation of radiation through absorbing, scattering medium, the RTE can be used, which takes the form:

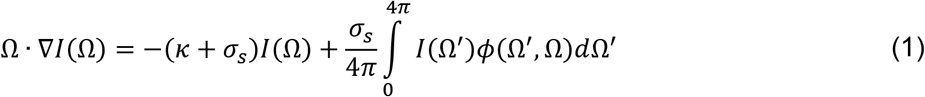

where Ω is the solid angle, *I* is intensity, *κ* is the absorption coefficient, *σ_s_* is the scattering coefficient, and *ϕ* is the scattering phase function.

Optical properties are highly wavelength-dependent. In this work we focus on values pertinent to blue light radiation, which has wavelengths of approximately 450 nm - 500 nm and is commonly used in mammalian light-inducible systems^[19–22]^.

The angular space in eq. 1 was discretized using the discrete ordinates method:

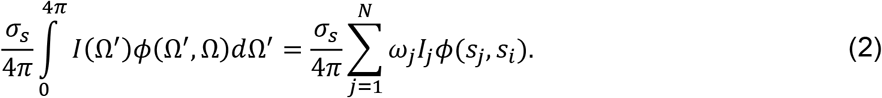

In eq. 2, the S6 level-symmetric even quadrature set and a linear discretization of the radiative intensity were used. Incident radiation was set to 7.958 W/m^2^·sr, which yields 100 W/m^2^, an order of magnitude of light intensities often used in optogenetics^[19–21,23]^. The radiation was emitted inwards from all boundaries of the domain.

### Calculation of absorption and scattering coefficients

The values of absorption and scattering coefficients used in this work are based on available data on mammalian cells in the literature^[24]^. Detailed calculations of parameters are available in Supporting Information. Given the high optical transparency of PBS, the mass absorption coefficient *α_a_* of mammalian cells is estimated from the absorption coefficient *κ* and the cell number density *ρ_c_*:

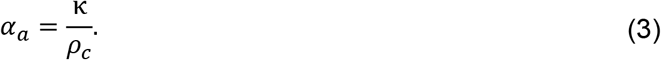

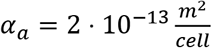 is used in this work, and this value of *α_a_* is used again in eq. 3 to calculate κ for various cell densities. Given culture medium also has a low absorption coefficient, calculations for the absorption coefficient as a function of cell density were also performed assuming negligible contributions from media^[25]^. An underlying assumption made for optically transparent media is the media is free of serum and phenol red, which is typically the case in industrial bioreactors.

For scattering, we use a reduced scattering coefficient of 2 m^-1^ and a scattering anisotropy *g* of 0.98^[26]^. The scattering coefficient was calculated as:

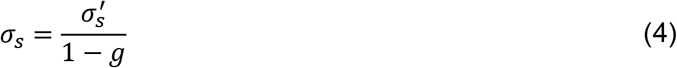

From eq. 4, the mass scattering coefficient *α_s_* is subsequently calculated:

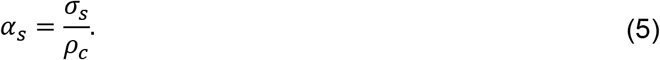

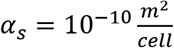 is used in this work. Similarly to *α_a_*, this *α_s_* value is reused in eq. 5 to calculate *σ_s_* for various cell densities. Scattering coefficients were also assumed to be unaffected by culture media^[25]^.

### Calculation of the scattering phase function

The scattering phase function was calculated using the Henyey-Greenstein function^[26,27]^:

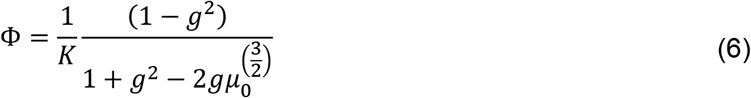

where *g* is the anisotropy factor, *μ*_0_ is the cosine of the scattering angle, and *K* is defined as the following to normalize the phase function:

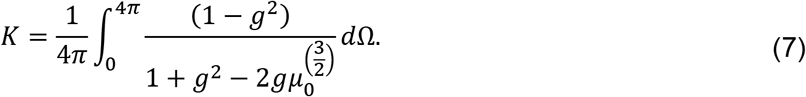

Eq. 6 and eq. 7 were substituted into eq. 1 to calculate radiative transfer.

## Results

To model light propagation through a bioreactor, we simulated a cylinder filled with medium, with light emitted inwards from the walls of the cylinder (Figure 1A). As light travels through the medium, which includes the cells, its intensity is decreased through absorption and scattering (Figure 1B).

**Figure 1.**
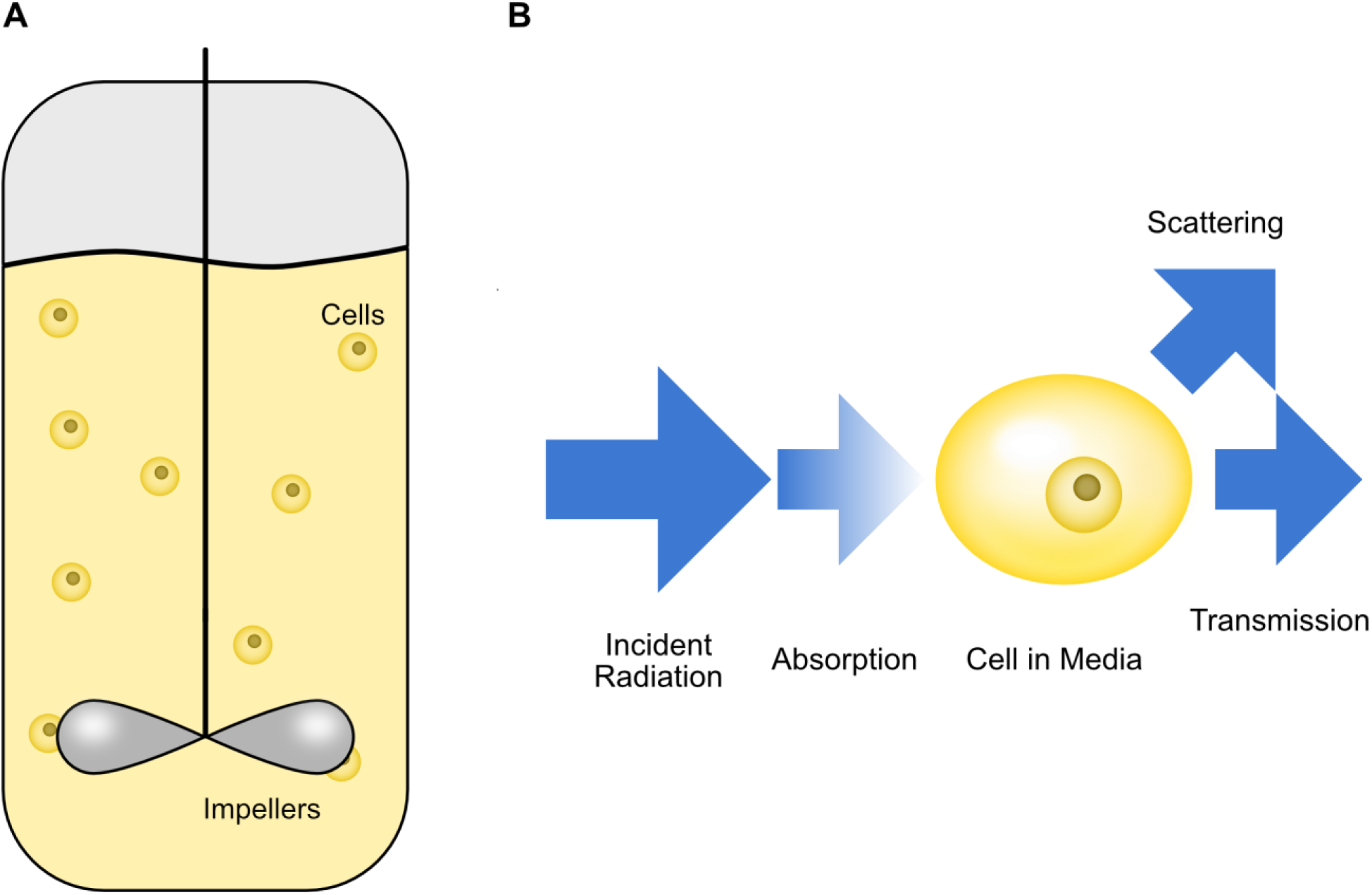
Light is attenuated through absorption and scattering as it travels through a bioreactor. (A) Cells dispersed in a well-mixed cylindrical bioreactor. (B) Radiation is absorbed, scattered, or transmitted while traveling through media containing cells.

First, we investigated the effects of the absorption coefficient, scattering coefficient, and scattering anisotropy on the intensity profile across a 100,000 L reactor, which is the order of magnitude size for the largest commercial bioreactors (Figure 2). We note that changing *g* also affects *σ_s_* when calculating from eq. 4, but for the purpose of observing the sensitivity of the intensity profile on *g, σ_s_* was held constant while scanning *g.* While scanning the values of one parameter, the others were kept at more transparent values such that the changes in intensity distribution from the parameter being scanned are more apparent. The range in values used for absorption and scattering coefficients were chosen such that they spanned values attained at minimum and maximum cell concentrations. Scattering anisotropy is known to be very high for mammalian cell suspensions, but lower values that tissues may have were also explored^[26–28]^.

**Figure 2.**
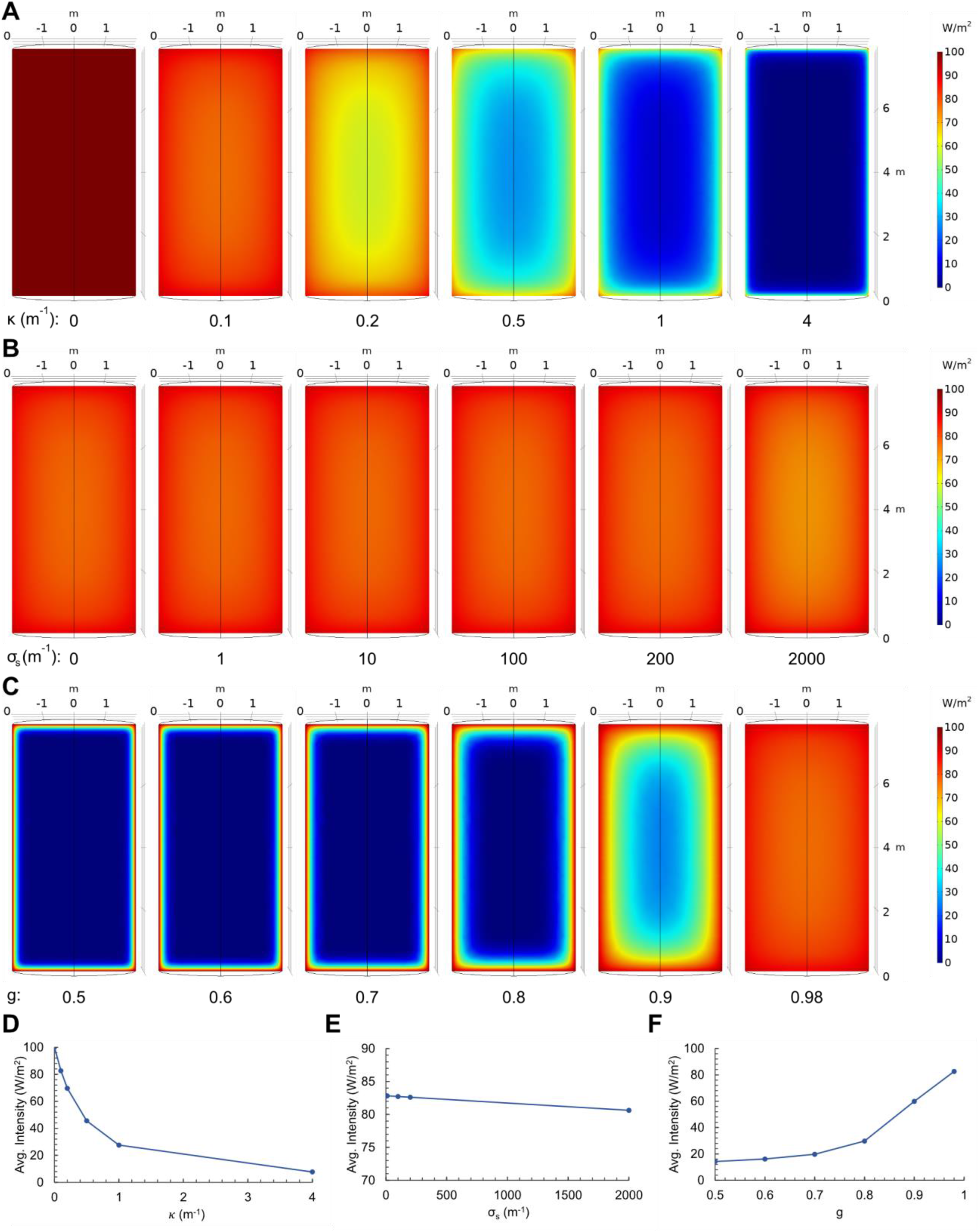
Light propagation over various *κ*, *σ_s_,* and *g.* (A) Parameter scan over κ with constant *σ_s_* and *g.* (B) Parameter scan over *σ_s_* with constant κ and *g.* (C) Parameter scan over *g* with constant κ and *σ_s_*. (D) Average intensity as a function of κ with constant *σ_s_* and *g.* (E) Average intensity as a function of *σ_s_* with constant κ and *g.* (F) Average intensity as a function of *g* with constant κ and *σ_s_.* When parameters were held constant, they were set to κ = 0.1 m^-1^, *σ_s_* = 200 m^-1^, and *g* = 0.98.

The parameter scan through relevant *κ* values indicated that absorption had a strong effect on light attenuation (Figure 2A). As *κ* increased, the intensity distribution shifted from mostly maximal throughout the reactor to mostly 0. In contrast, increases in *σ_s_* did not noticeably impact light propagation until the maximum value of 2000 m^-1^ was used (Figure 2B). Although scattering coefficients were high, the high scattering anisotropy may alleviate decreases in intensity along the radial direction by scattering the radiation strongly in the forward direction. Indeed, scanning lower scattering anisotropy values caused lower intensities in the center of the reactor by redirecting the radiation away from the radial path (Figure 2C). This indicates that *σ_s_* was not negligible, even at the intermediate value of 200 m^-1^ that was used in the anisotropy sweep, as changing anisotropy would not impact the intensity distribution if *σ_s_* was too low. Rather, strong scattering occurred but simply in the forward direction, and decreasing the scattering anisotropy had a very strong effect on the light intensity distribution. The change in intensity distribution from changing the scattering anisotropy is only attributed to scattering, and not absorption.

Mammalian cell bioreactors are utilized at a wide range of sizes and aspect ratios^[14–16]^. The surface area to volume ratio of a cylinder decreases as 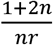, where *n* is the height to radius aspect ratio and *r* is the radius. For this system, in which the light is emitted inwards from the reactor walls, the average intensity of light in the reactor is expected to decrease as the reactor size increases. We first observed the effect of reactor size and height-to-diameter (HD) ratio on intensity, using the maximum cell density 2 · 10^13^ cells/m^3^ and absorption and scattering coefficients at that cell density (Figure 3).

**Figure 3.**
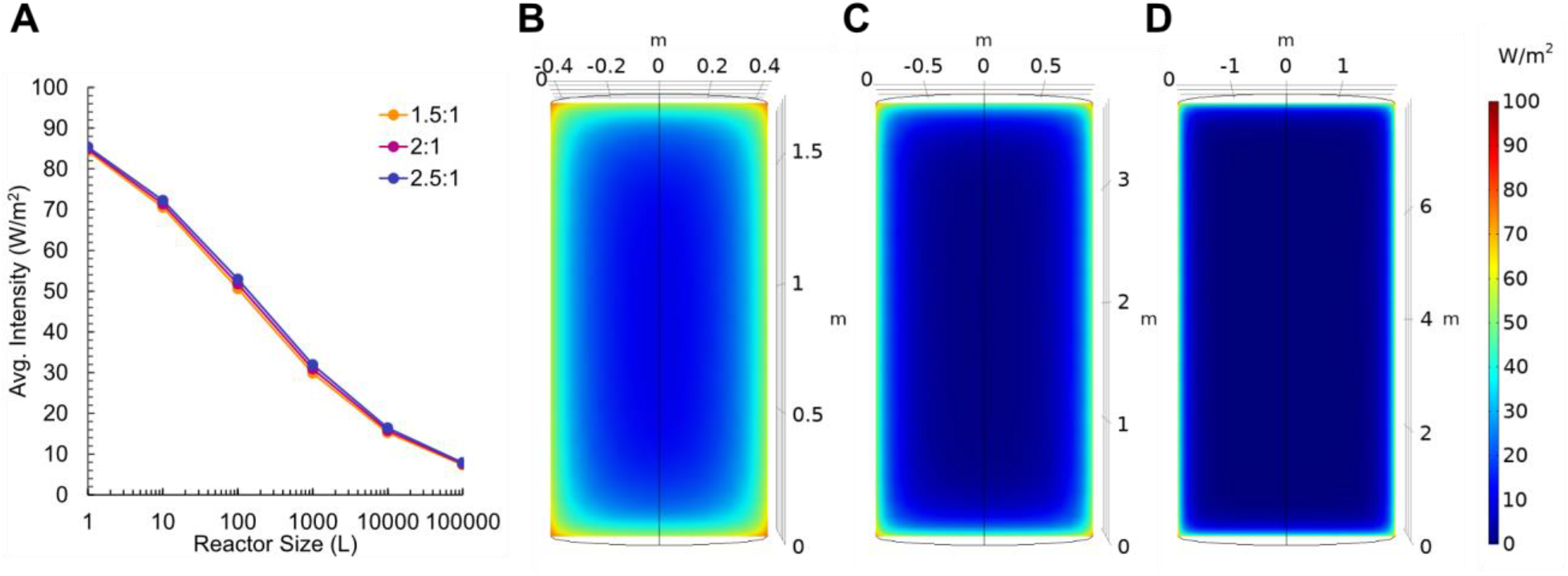
Light propagation vs HD ratios and volume. (A) Average intensity in the reactor as a function of reactor volume for three different HD ratios. (B-D) Slices of the 3D radial intensity distributions for (B) 1,000, (C) 10,000, and (D) 100,000 L reactors.

As reactor volume increased, the volume average intensity decreased. The HD ratio also affected the average intensity, where larger HD ratios, which yield a higher surface area to volume ratio, had higher average intensity, as expected. Overall, however, the aspect ratios in the range of interest have little effect on the average intensity in the relevant range for mammalian cell culture.

As a metric for sufficient light-activation, we used the light intensity at which 90% of maximal expression was obtained in previous work, which corresponds to 50 W/m^2[19]^. For the 1,000 L and larger bioreactors, the average intensity was below the threshold of 50 W/m^2^, which indicates that there may be insufficient activation. For these cases, we inspected slices of the 3D radial intensity distributions for the bioreactor with 2:1 HD ratio (Fig. 3B-3D). For the larger reactors, most reactor volumes had near-zero intensity, but positions near the boundaries still had high intensity. Under this constraint, we now consider that in many optogenetic applications^[13,19,21]^, cells are not required to be constantly illuminated, but are sufficiently activated following short pulses of light followed by extended periods in the dark. Short millisecond to second pulses of light followed by seconds or minutes in the dark are sufficient for activation because the timescale for the optogenetic proteins to deactivate through conformational changes and dissociation are longer, with half-lives on the order of seconds to minutes^[29]^. Therefore, reactors that have a lower average intensity than the required intensity may still be able to activate cells if regions near the boundary meeting the threshold intensity are sufficiently large. As an order-of-magnitude estimate, we considered a case in which 1% of time spent under illumination is sufficient for activation^[29]^. In the context of a bioreactor with constant illumination, if a cell occupies a region of the reactor with the threshold intensity for 1% of the time, the cell is adequately activated. Then, assuming a well-mixed system, the volume fraction of the reactor achieving threshold intensity is equal to the time-averaged probability that a cell will experience that intensity. Thus, for the larger bioreactors, reactor volume fractions achieving threshold intensity were calculated (Fig. 4A). The 2:1 HD ratio is considered as a representative case, and the other aspect ratios are expected to yield similar results as the intensity distributions were similar (Fig. 3A).

**Figure 4.**
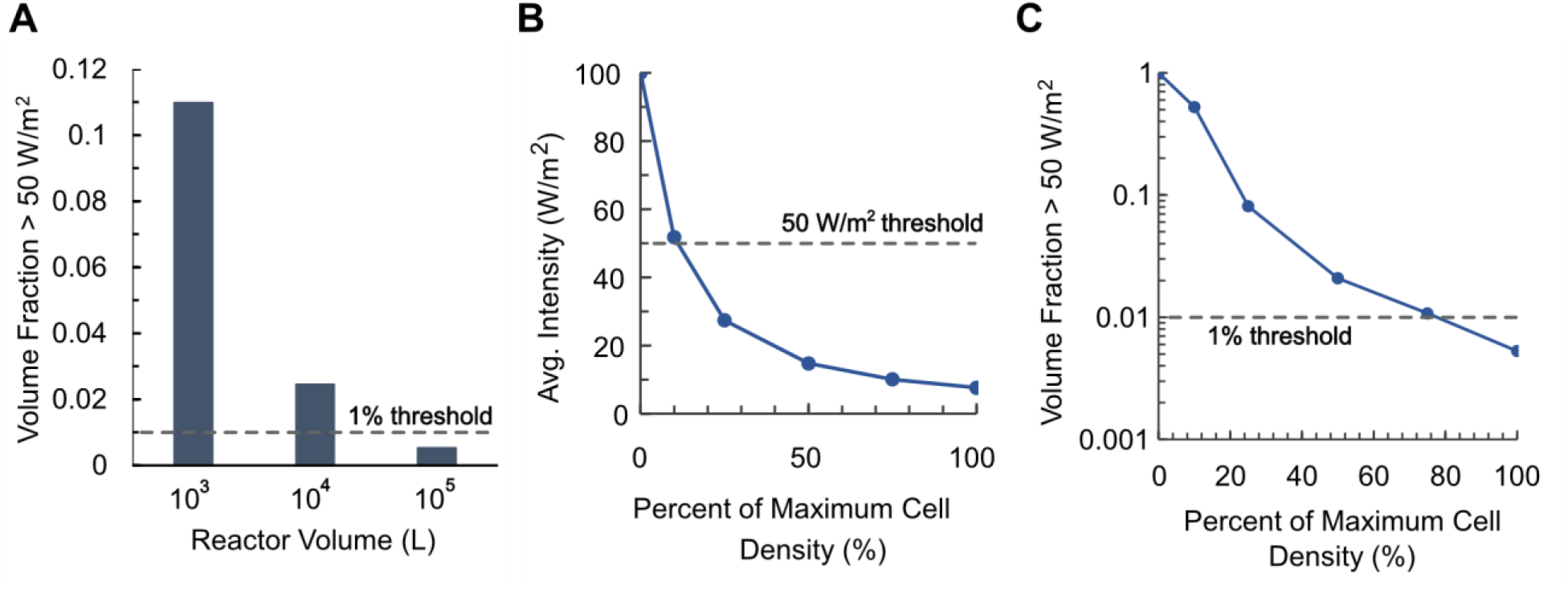
Volume fractions achieving threshold intensity for 2:1 HD ratio reactors. (A) Volume fractions achieving threshold intensity at maximum cell density. (B) Average intensity as a function of cell density. (C) Volume fractions achieving threshold intensity as a function of cell density. Maximum cell density was taken to be 2·10^7^ cells/mL. Dashed lines indicates threshold volume fraction and threshold average intensity for a well-mixed system.

The 1,000 L and 10,000 L reactors now met the threshold for adequate activation, as more than 1% of the reactor volumes had intensities > 50 W/m^2^. The 100,000 L bioreactor did not meet the volume fraction threshold. Thus, for the largest of industrial bioreactors with the highest cell densities, there was a decrease in the dynamic range for light activation. Linearly interpolating as a first approximation, the largest reactor size that met the volume fraction threshold for light activation with maximum cell culture density occurred at approximately 80,000 L for the parameters tested.

Finally, we explored the effects of increasing cell density. As cell density increases, absorption and scattering increase proportionally, as given in eq. 3 and eq. 5. Considering the range of cell density from 0 cells/mL – 2·10^7^ cells/mL, the effects of cell growth on the intensity profile in the 100,000 L bioreactor were explored through simultaneous increases in *κ* and *σ_s_.* As cell density was increased, *κ* and *σ_s_* were recalculated from *α_a_* and *α_s_* and the average intensities inside the reactor were calculated (Fig. 4B). From the parameter scan, we determined that the highest density of cells that can be sufficiently activated in a 100,000 L bioreactor is approximately 75% of maximum density, or 1.5·10^7^ cells/mL (Fig. 4C).

## Discussion

Our data suggest that light-inducible systems are viable in large-scale mammalian suspension culture bioreactors. The average intensity drops quickly as reactor size or cell density increases, but only short pulses of light are required for robust light-activation. This allows the high intensities near the boundaries of the reactor to sufficiently activate cells, even when the average intensity within the overall volume is low.

In general, absorption dramatically limits light propagation as the absorption coefficient or cell density increases. However, scattering does not have as large an effect, even though scattering coefficients of mammalian cells are very high. The lower effect of scattering on the intensity distribution is due to the scattering anisotropy also being high. High scattering anisotropy causes most of the radiation to be forward scattered, mitigating the attenuation of radiation in the radial direction of the reactor. The effects of scattering became more evident when the scattering anisotropy was decreased, which rapidly decreased the intensity throughout the volume (Fig. 2C). Sparging was not considered in this work, but the presence of bubbles will increase scattering. Scattering from bubbles will require careful and more intensive computation, as the extent of scattering depends on the bubble distribution throughout the reactor, which may not be as uniformly dispersed as the cells. The bubble size distribution also affects scattering, where scattering is affected by the width of the size distribution and cannot be calculated solely from the mean bubble size^[26,30]^. The sparging rate will also affect the size and total bubble volume in the reactor.

The maximum reactor size that can support optogenetics based on the geometry and parameters used in this work is approximately 80,000 L, but other elements that may be present in a real reactor may be utilized to further enable optogenetics. For instance, bioreactors are often equipped with impellers, and LEDs may be mounted on the impeller shaft, such that illumination occurs from the center of the reactor, as well as from the reactor walls. Baffles may also be added to the geometry, which increases the surface area available for LED integration without increasing the volume. Finally, LED intensities can be increased, though care must be taken to avoid phototoxicity to the cells^[31]^.

Simulations in this work explore maximal mammalian bioreactor sizes, which are several orders of magnitudes larger than photobioreactors for algae and cyanobacteria ^[7,9,10]^. Larger reactors can be supported by optogenetics because sufficient activation only requires illumination comprising a small fraction of overall time, such as 1% (Fig. 4). We speculate that lower illumination fractions are sufficient for optogenetics because less photon energy is required to saturate single gene circuits or signaling pathways, whereas light-dependent cell growth may require more energy in algal and cyanobacterial systems.

The extent to which light propagates through mammalian suspension cultures heavily depends on the wavelength of the incident radiation. The simulations performed in this work use parameter values assuming blue light at approximately 450 nm. Blue light optogenetics systems are the most widely used, but systems utilizing other wavelengths are also gaining popularity. In particular, red light optogenetics is promising because mammalian cells have lower absorption and scattering for red light, which would increase the penetration of red light through larger systems ^[24,26]^. Absorption and scattering generally decrease as the wavelength increase to the infrared region, and the results in this work may be interpreted as conclusions from conservative calculations. As the use of higher wavelength optogenetics systems increases, further scaleup will become feasible in mammalian suspension cultures or enable deeper penetration of light into tissues, such as in cultivated meat applications.

## Conclusion

We demonstrate that light propagates sufficiently in large bioreactors of mammalian suspension cultures. While we did observe the limits of optogenetic activation in the largest scales of industrial bioreactors, modifications to the lighting conditions and bioreactor geometries may extend the limits of optogenetics into volumes of higher orders of magnitudes. Future efforts to model more realistic mammalian bioreactor setups, including sparging, impellers, and baffles, will improve the accuracy of results. Ultimately, such simulations can be used to inform new photobioreactor design in growing industries requiring innovation to lower production costs of complex products.

## Acknowledgments

We would like to thank Dr. Adele Igel for her feedback on specific technical aspects of this manuscript. This work was supported by a Hellman Fellowship to PSS and a summer research fellowship from the University of California, Davis College of Engineering to SAM.

