## Supporting Information for "Computational evaluation of light propagation in cylindrical bioreactors for optogenetic mammalian cell cultures"

### Calculation of mass absorption and scattering coefficients

With an absorption coefficient of 0.04 $cm^{-1}$ at 60 million cells suspended in 3 mL phosphate buffered saline (PBS) at 450 nm is approximately 0.04 $cm^{-1}$[1]. Given the high optical transparency of PBS, the mass absorption coefficient $\alpha_a$ of mammalian cells is estimated from the absorption coefficient $\kappa$ and the cell number density $\rho_c$:

$$\alpha_a = \frac{\kappa}{\rho_c}.$$

Substituting values, we find

$$\alpha_a = \left(\frac{0.04}{cm}\right)\left(\frac{3cm^3}{(6 \cdot 10^7 cells)}\right)\left(\frac{m^2}{100^2 cm^2}\right) = 2 \cdot 10^{-13}\ \frac{m^2}{cell}.$$

For scattering, we use a reduced scattering coefficient of 2 $cm^{-1}$ and a scattering anisotropy $g$ of 0.98[2]. The scattering coefficient was calculated as:

$$\sigma_s = \frac{\sigma_s'}{1-g}.$$

The scattering coefficient is calculated to be:

$$\sigma_s = \left(\frac{2cm^{-1}}{1-0.98}\right)\left(\frac{100cm}{m}\right) = 10^4 m^{-1}.$$

Subsequently the mass scattering coefficient $\alpha_s$ is calculated as:

$$\alpha_s = \frac{\sigma_s}{\rho_c}$$

Substituting:

$$\alpha_s = \frac{10^4 m^{-1}}{10^8 \frac{cell}{mL}}\left(\frac{m^3}{10^6 mL}\right) = 10^{-10}\,\frac{m^2}{cell}.$$

**<u>Height-to-diameter ratio of a cylinder</u>**

For a cylinder with a height-to-diameter (HD) ratio of n, the surface area to volume ratio can be expressed as:

$$\frac{2\pi r^2 + 4\pi n r^2}{2\pi n r^3}$$

where $n$ is the HD ratio.

This can be simplified to:

$$\frac{1 + 2n}{nr}.$$
